# Evolution of immune infiltration and immune triad formation across early melanoma progression

**DOI:** 10.64898/2026.10.29.762853

**Authors:** Kayla A Marks, Abigail Florell-Orton, Min Hu, Burke Lawlor, Scott Florell, Mitchell S Stark, Kathleen J Green, Dekker C Deacon, Frederick R Adler, Robert L Judson-Torres, Melissa Q Reeves

## Abstract

Tumor-infiltrating lymphocytes (TIL) in melanoma are a clinically important prognostic marker and are routinely used to inform patient care. However, when immune recognition of transformed melanocytes begins and how it evolves during early melanoma progression is not well understood. Using spatial transcriptomics and a unique cohort of patient-matched lesions representing multiple stages of early evolution, we investigate T cells, dendritic cells (DCs) and multi-cell immune “triads” during melanoma initiation. We find that T cell and DC density, triad density, and triad proximity to melanoma cells are elevated in regions of Stage I invasive melanoma as compared to Stage 0 in situ lesions and nevi. Further, we find that triad abundance correlates with both T cell and DC density, but is not predicted by clustering patterns of these cell types over larger spatial scales. We conclude that immune recognition of melanoma and engagement with the lesion significantly increases with progression from Stage 0 to Stage I disease. Further, using nevi that are depth-matched in the dermis to Stage I invasive lesions, we demonstrate this increased immune recognition is not a consequence of dermal invasion alone. This points to a paradigm where immune engagement of invasive melanoma reflects recognition of melanoma cell-intrinsic changes that emerge in Stage I disease.

**Significance:** Tumor-infiltrating lymphocytes predict better melanoma patient outcomes even in early stage disease, but how and when immune recognition of melanoma is initiated remains poorly understood. Using patient-matched lesions of multiple progression stages, we find immune engagement with melanoma strongly increases between Stage 0 in situ and Stage I invasive disease.

## Introduction

Melanoma is an aggressive form of skin cancer whose microenvironment contains immune cell interactions that can either promote anti-tumor responses or drive immune evasion. The responsiveness of melanoma to immune control has led to the widespread adoption of immunotherapies to treat patients with invasive or metastatic disease (1). T cell infiltration into melanoma has been recognized for decades, with tumor-infiltrating lymphocyte (TIL) scoring used both as a positive prognostic marker and as a predictor of immunotherapy efficacy even in early stage tumors (2–11).

Early melanoma can progress through defined histological stages: nevi, in situ, and invasive. Melanocytic nevi are benign tumors of melanocytes. In situ melanoma (Stage 0) contains neoplastic cells that remain contained within the epidermis and have no detectable cells present in the dermis that would indicate invasive melanoma (12). Both nevi and in situ disease are potential precursor lesions to invasive melanoma (stages I-II) – defined as melanoma which has invaded the deeper layer of skin, the dermis (13,14). The prognostic value of TIL has been most widely evaluated in invasive melanoma, and higher TIL grades in stage I-II invasive melanoma are positively associated with patient survival and with decreased risk of positive sentinel lymph nodes (2,3,5,8,11). Two TIL scoring systems are routinely used: the Clark system, which divides TIL into “brisk” (lymphocytes infiltrating the entire base of the tumor), “non-brisk” (focal or patch lymphocytes), and “absent”, and a scale promoted by the Melanoma Institute of Australia (MIA) which divides TIL instead into 4 categories with finer gradation on the abundance of TIL foci (8,15). Notably, while these scales are routinely used for invasive melanoma, comparable scoring of TIL in nevi and in situ melanoma is far less frequently performed, and the evolution of TIL patterns from these precursor lesions to invasive melanoma has been described in only a handful of studies, with the focus primarily on quantifying T cells (15,16).

While T cells are critical for effective tumor control and the basis of nearly all clinically utilized immunotherapies in melanoma, T cells cannot act alone. Activation and expansion of cytotoxic CD8+ T cells requires interaction with antigen-presenting cells, and in tumor contexts, is specifically dependent on dendritic cells (DCs) (17,18). T cell-mediated rejection of melanomas has been shown to depend on physical co-localization of CD8+ T cells and DCs in and around the tumor (19,20). Further, the importance of DC-T cell interactions at the tumor site has been underscored by the recent discovery of “triads” – functional immune units comprised of a DC, CD8+ T cell and CD4+ T cell – which are critical for effective tumor control and whose presence has been shown to be more important in predicting immune control and response to immunotherapy than the abundance of T cells or DCs alone (21–23).

Here, we make use of a spatial transcriptional profiling dataset to interrogate the abundance and spatial organization of multiple T cell and dendritic cell subsets in nevi, in situ, and locally invasive melanoma. As a unique feature of this dataset, multiple stages of early disease were present in distinct regions of the same tissue specimen. These regions were annotated by a dermatopathologist, enabling us to examine immune cell patterns associated with different stages of disease from within the same patient. To robustly quantify immune infiltration patterns, we measured T cell and DC density, T cell and DC clustering, and formation of DC-CD4+ T cell-CD8+ T cell “triads” as independent methods for evaluating immune engagement with melanomas at each stage of progression. We find that density of both T cells and DCs is markedly increased in invasive disease as compared to both in situ lesions and nevi, and these cell types exhibit non-random clustering in nearly all melanoma regions where they are present. Strikingly, we additionally find that DC-CD4+ T cell-CD8+ T cell “triads” are both more abundant in invasive lesions and, when outside a lesion boundary, are significantly closer to invasive tumor regions than to melanoma in situ or nevi, implying that immune engagement is dramatically increased in and around invasive melanoma. While we find multiple DC subsets engaged in these triads, we find that Langerhans cells, DC-like skin resident antigen presenting cells, rarely participate. Together, our data reveal that immune engagement with melanoma, as quantified by multiple axes, significantly increases specifically between Stage 0 in situ and Stage I invasive disease.

## Results

### T cell density increases from in situ to invasive disease

We assembled a cohort of melanomas with dermatopathologist-annotated regions of nevus, in situ, and invasive disease within the same tissue samples (**Fig. 1A**). Our cohort comprised a total of 15 specimens from 13 patients, within which we annotated 12 nevi, 40 regions of in situ disease, and 29 invasive regions. Clinical notes indicated the presence of non-brisk TIL in 12 of the 13 patients, while TIL were annotated as absent in the remaining case. Spatial transcriptional analysis using the 10X Xenium platform identified all expected cell types in these specimens, including tumor cells, immune cells, stromal cells and keratinocytes (**Supp. Fig. 1**). Given our interest in TIL patterns, we next focused on creating high-quality spatial cell maps of T cells and dendritic cells (DC) and their subsets (**Fig. 1B-D**). We analyzed the spatial patterns of these critical immune cell players across early progression stages. T cells were divided into CD8+ T cells, non-regulatory CD4+ T cells (hereafter “CD4+”), and regulatory CD4+ T cells (hereafter “Treg”; defined by expression of FOXP3). DCs were divided into CLEC9A+, CLEC10A+ and LAMP3+ DCs. *CLEC9A* and *CLEC10A* mark conventional type 1 and type 2 DCs (cDC1s and cDC2s) respectively, while *LAMP3* marks activated DCs (24).

**Figure 1.**
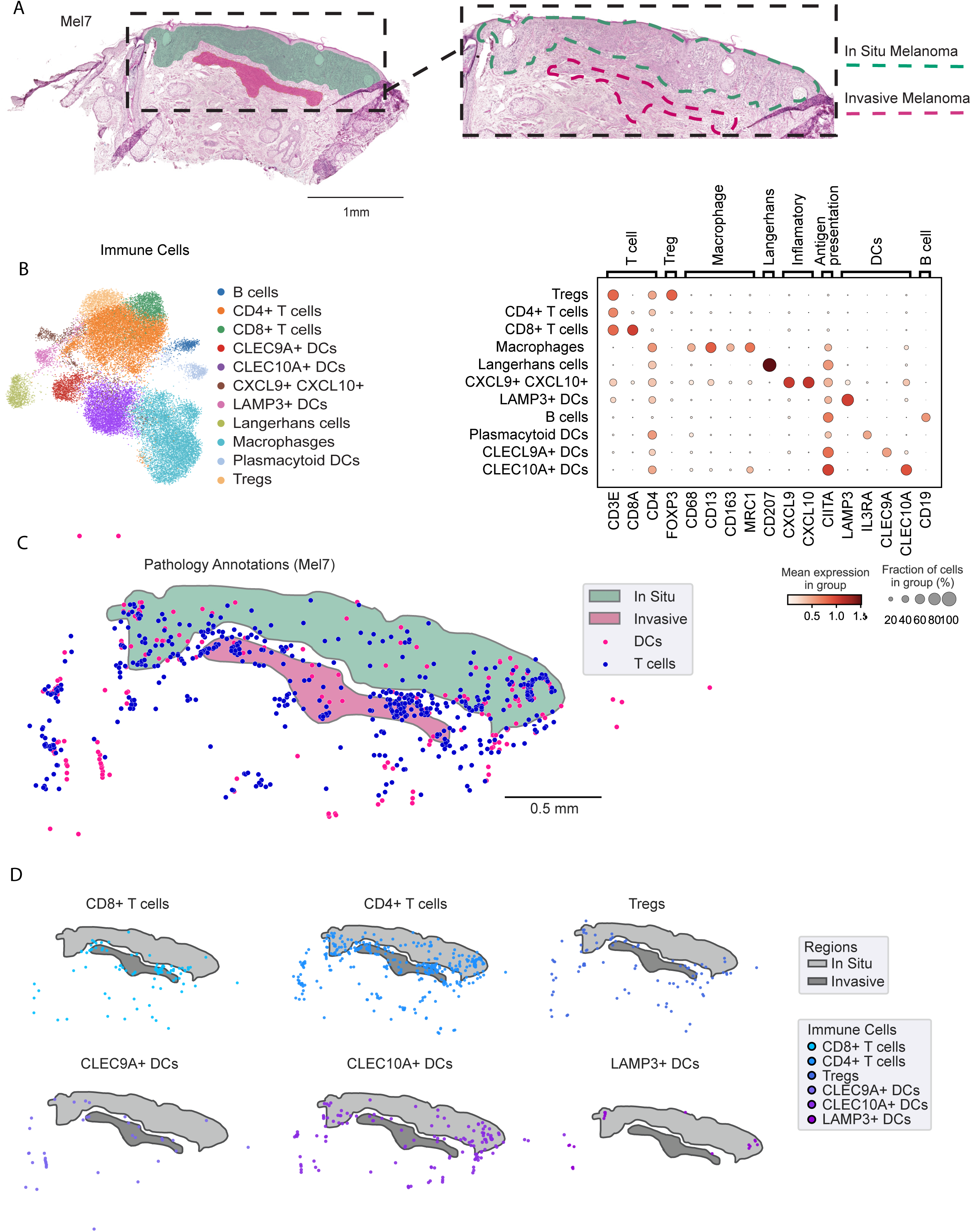
Spatial maps of immune cell subsets in early melanoma. Immune cell identities, spatial positions of immune cells, and regions of nevi, in situ melanoma and invasive melanoma were identified in a Xenium spatial transcriptomics dataset comprised of 15 samples from 13 patients. (A) H&E of a representative sample with dermatopathologist-annotated regions of in situ and invasive disease, annotated in green and pink respectively. (B) UMAP (left) of immune cell subsets identified across all samples and dot plot (right) showing expression of lineage marker genes. (C) Overlay of dendritic cells (DCs, pink) and T cells (blue) on pathologist-annotated regions from a representative sample. (D) The same lesion regions as (C), showing individual T cell subsets (CD4+ T cells, CD8+ T cells, and regulatory T cells (Tregs)) and DC subsets CLEC9A+ cDC1s, CLEC10A+ cDC2s, and LAMP3+ activated DCs.

Recapitulating prior studies (16), we found T cell density increases from in situ to invasive disease (**Fig. 2A**). Total T cell density was significantly higher in invasive disease (p < 0.0001, Mann-Whitney non-parametric), as was the density of CD4+ T cells, CD8+ T cells, and regulatory T cells (all p < 0.0001). Total DC density was also higher in invasive disease (**Fig. 2B**, p = 0.002), an effect that was most pronounced among CLEC9A+ and LAMP3+ subsets that have been previously implicated as critical players in anti-tumor immunity (22,25) (CLEC9A+, p < 0.0001; LAMP3+, p = 0.0011; CLEC10A+, p = 0.0122). To complement these region-level comparisons, we modeled cell counts against the fractional area of each lesion type across a uniform grid and computed odds ratios for immune cell presence in invasive versus in situ disease (**Fig. 2C-E**). T cells had higher odds of being in invasive regions in 11 of 12 melanoma samples (**Fig. 2D**). DCs showed a weaker, more variable association, with higher odds in invasive regions in only 4 samples, higher odds in in situ regions in 2 samples, and no association in 6 samples, indicating that DC enrichment in invasive disease is sample-dependent (**Fig 2E**).

**Figure 2.**
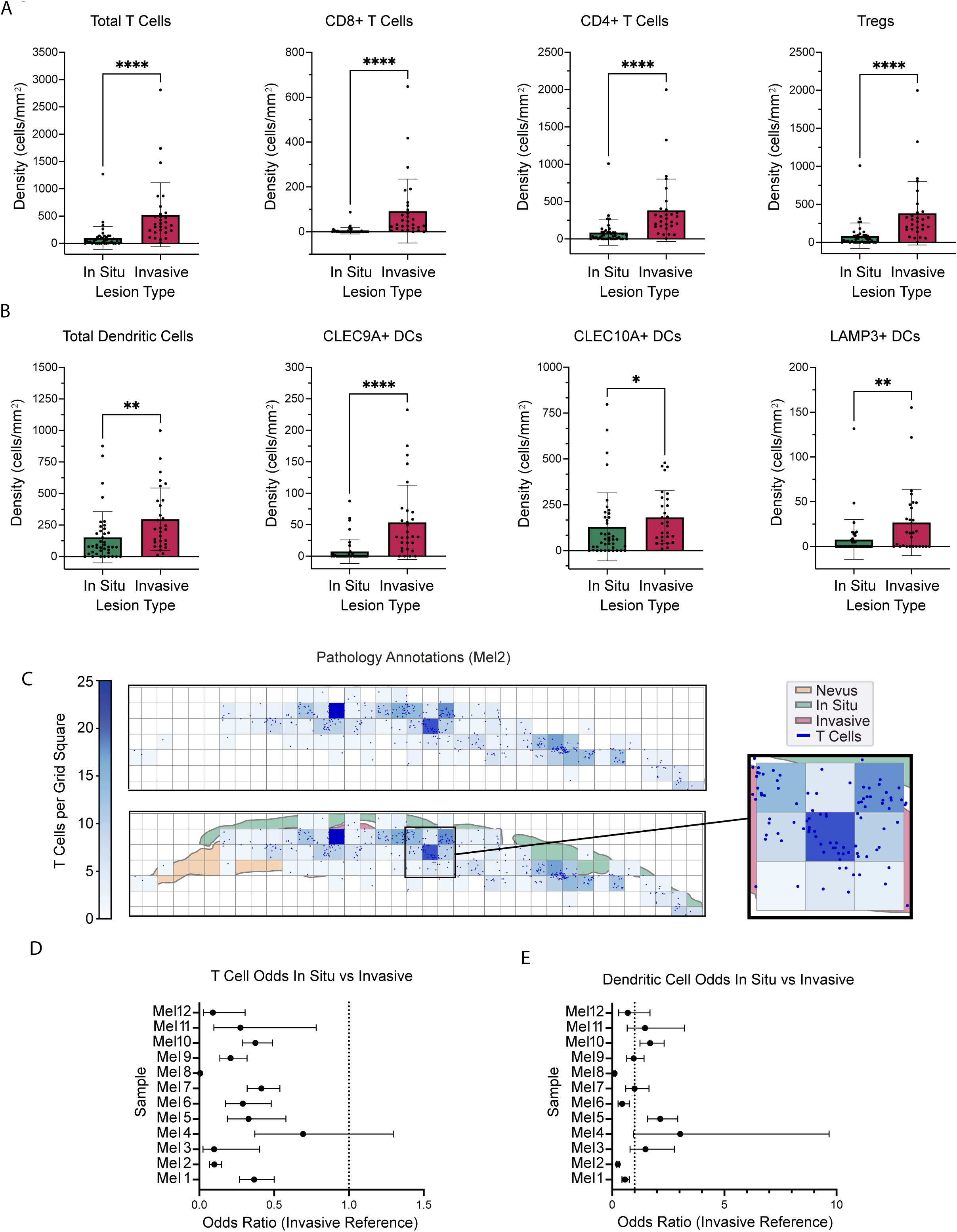
T cells and DCs are more abundant in invasive than in situ disease. (A, B) Bar graphs showing density of total T cells, CD8+ T cells, CD4+ T cells and Tregs (A) and total dendritic cells (DCs), CLEC9A+ DCs, CLEC10A+ DCs, and LAMP3+ DCs (B). Statistical significance in (A-B) was evaluated by Mann-Whitney test. \**p*<0.05, \*\**p*<0.01, \*\*\*\**p*<0.0001. (C) Schematic of grid-based analysis. Squares were drawn over the entire sample and for each square, the fractional area belonging to nevus regions, in situ regions, invasive regions, or none of the above was calculated, as was the total number of T cells or DCs in the square. These values were used for linear regression and odds ratios analyses. The figure shows a representative region with the pathology annotations (orange, green, and pink) overlaid with a heatmap showing T cell density (blue scale) for squares with ≥1 T cell and exact T cell positions in dark blue. (D, E) Odds ratios (OR) plots showing odds of T cells (D) and DCs (E) being in regions of in situ versus invasive disease. Invasive disease is used as the reference such that an OR > 1 indicates higher odds of the cells being in an in situ region and an OR < 1 indicates higher odds of the cells being in invasive regions. Error bars show 95 percent confidence interval.

### T cell enrichment tracks malignancy, not dermal invasion

Invasive disease is, by definition, deeper in the skin than in situ disease, having invaded from the epidermal home of melanocytes into the dermis. This raises the possibility that the density-invasion association reflects easier immune cell access to deeper tissue or an immunologic response to melanocytes entering a foreign tissue, rather than malignant transformation itself. Nevi are benign yet can also reside in the dermis, often at similar or even greater depth than the invasive disease. We exploited this feature to decouple dermal depth from malignancy and confirmed that the depth of nevi and of invasive regions was not different in our cohort across the 6 samples that included both nevi and invasive regions (**Fig. 3A, B**). Despite being at similar depth, total T cell density was significantly higher in invasive disease than in nevi (p = 0.0007), as was total DC density (p = 0.0006) (**Fig. 3C-D**). Among subtypes, the effect was strongest for CD4+ (p = 0.002) and CD8+ T cells (p = 0.004) and not significant for Tregs, although Tregs also trended toward an increase in invasive regions (**Fig. 3C**). DC subtypes also showed significant differences (CLEC9A+, p = 0.003; LAMP3+, p = 0.044; CLEC10A+, p = 0.019) (**Fig. 3D**). Odds-ratio analysis was consistent, favoring invasive regions in 3 of 6 specimens for T cells (**Fig. 3E**) and 4 of 6 specimens for DCs (**Fig. 3F**). Thus, immune cells infiltrate invasive disease more than depth-matched nevi, indicating that T cell and DC enrichment segregates with malignant progression rather than dermal location.

**Figure 3.**
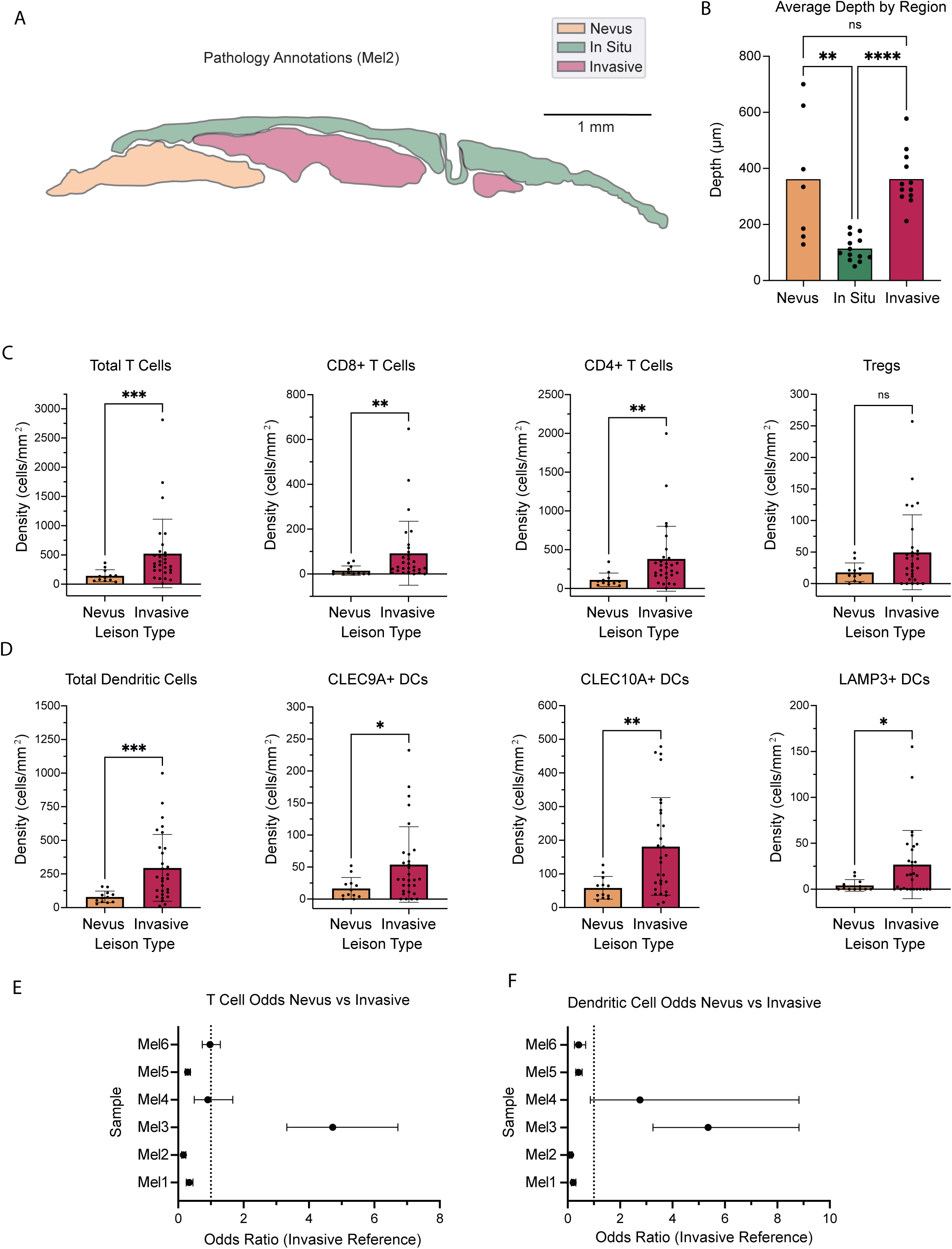
T cells and DCs are more abundant in invasive melanoma than depth-matched nevi. (A) Pathology annotation of a representative sample including nevus, in situ, and invasive regions of melanoma. (B) Bar graph showing depth from the granular zone of nevus, in situ, and invasive regions across our cohort. Statistical significance based on Kruskal-Wallis test. (C, D) Bar graphs showing density of total T cells, CD4+ T cells, CD8+ T cells and Tregs (C) and total DCs, CLEC9A+ DCs, CLEC10A+ DCs, and LAMP3+ DCs (D). Statistical significance in (C-D) was evaluated by Mann-Whitney test. (E-F) Odds ratio (OR) plots showing odds of T cells (E) and DCs (F) being in regions of nevus vs invasive disease. Invasive disease is used as the reference, such that an OR > 1 indicates higher odds of the cells being in nevus region and an OR < 1 indicates higher odds of the cells being in invasive regions. Error bars show 95 percent confidence interval. \**p*<0.05, \*\**p*<0.01, \*\*\**p*<0.001, \*\*\*\**p*<0.0001.

### T cells and DCs exhibit clustered spatial patterns within melanoma regions

Measuring density alone fails to capture the spatial organization of cells within a region. As the spatial organization of immune cells within a tumor has emerged as an important prognostic factor for patients, we next asked whether T cells were clustered with one another within early melanoma lesions. To quantify clustering, we implemented Ripley’s K with L(r) – r transformation and measured the integral of this function from 7 to 35μm (**Supp. Fig. 2A**), representing a spatial scale of approximately one to five T cell-diameters, and named this metric the Ripley’s K Clustering (RKC) score. Ripley’s K normalizes neighbor counts against the global point density to produce a measure of clustering that is independent of density, and RKC score values greater than zero indicate clustering, while values below zero indicate dispersion (e.g., a preference for cells to be further away from a neighbor than they would be at random) (**Supp. Fig. 2B**). The significance of the RKC score was computed by Monte Carlo simulation. We constrained our analysis to regions with at least 30 total T cells (26,27) and where the shortest dimension of the region was >140μm (4 times the upper limit of our integral) (28,29). Using these constraints, we were able to evaluate clustering in 6 of 12 (50%) nevus regions, 6 of 40 in situ regions (15%), and 19 of 29 (66%) invasive regions. Of note, the area of nevi tended to be larger than in situ and invasive regions, and this resulted in a larger proportion of nevi than in situ regions meeting the 30 T cell requirement for evaluation despite exhibiting similarly low T cell density. We found that T cells exhibited significant clustering (RKC > 0, *p* < 0.05) for all regions analyzed (**Supp. Fig. 2C**). We evaluated clustering of DCs using a similar approach and observed that DCs exhibited significant clustering in one evaluable nevus (all other nevi contained <30 DCs), in 10 of 10 evaluable in situ regions, and in 11 of 12 evaluable invasive regions (**Supp. Fig. 2D**). As such, both T cells and DCs exhibit non-random, clustered patterns of infiltration when present in early melanomas of all stages.

### T cell-DC triads are associated with melanoma progression

T cell-DC triads, in which a DC simultaneously interacts with a CD8+ and a CD4+ T cell, have been identified as critical hubs of immune activity in solid tumors, and their presence in patients with advanced disease is associated with better responses to immune checkpoint inhibitor therapy (21,22). We defined immune triads in our dataset as being comprised of a DC (of any subset), a CD4+ T cell (non-Treg) and a CD8+ T cell all within 10.6 um of one another (∼1.5 T cell diameters) (**Fig 4A**). Interestingly, cells annotated as “CXCL9+ CXCL10+” (**Fig. 1B**), which contained a mixture of dendritic cell and T cell markers, were found in triad structures with CD4+ and CD8+ T cells. Dendritic cells engaged with T cells have been reported to be strong producers of CXCL9 and CXCL10 (19,30) which attract T cells through the CXCR3 receptor. The participation of these cells in T cell interactions explains the presence of *CD3E* transcripts that had previously prevented us from confidently classifying them as DCs. Based on these observations, we included in this analysis triads comprised of a CXCL9+ CXCL10+ DC, CD4+ T cell and CD8+ T cell. Strikingly, we found that triads were significantly more abundant in invasive regions as compared to nevi (**Fig. 4B**). These results indicate that the increase in T cells and DCs in invasive melanoma noted above is more than a stochastic accumulation of immune cells, but rather coincides with the formation of active immune hubs in response to invasive disease.

**Figure 4.**
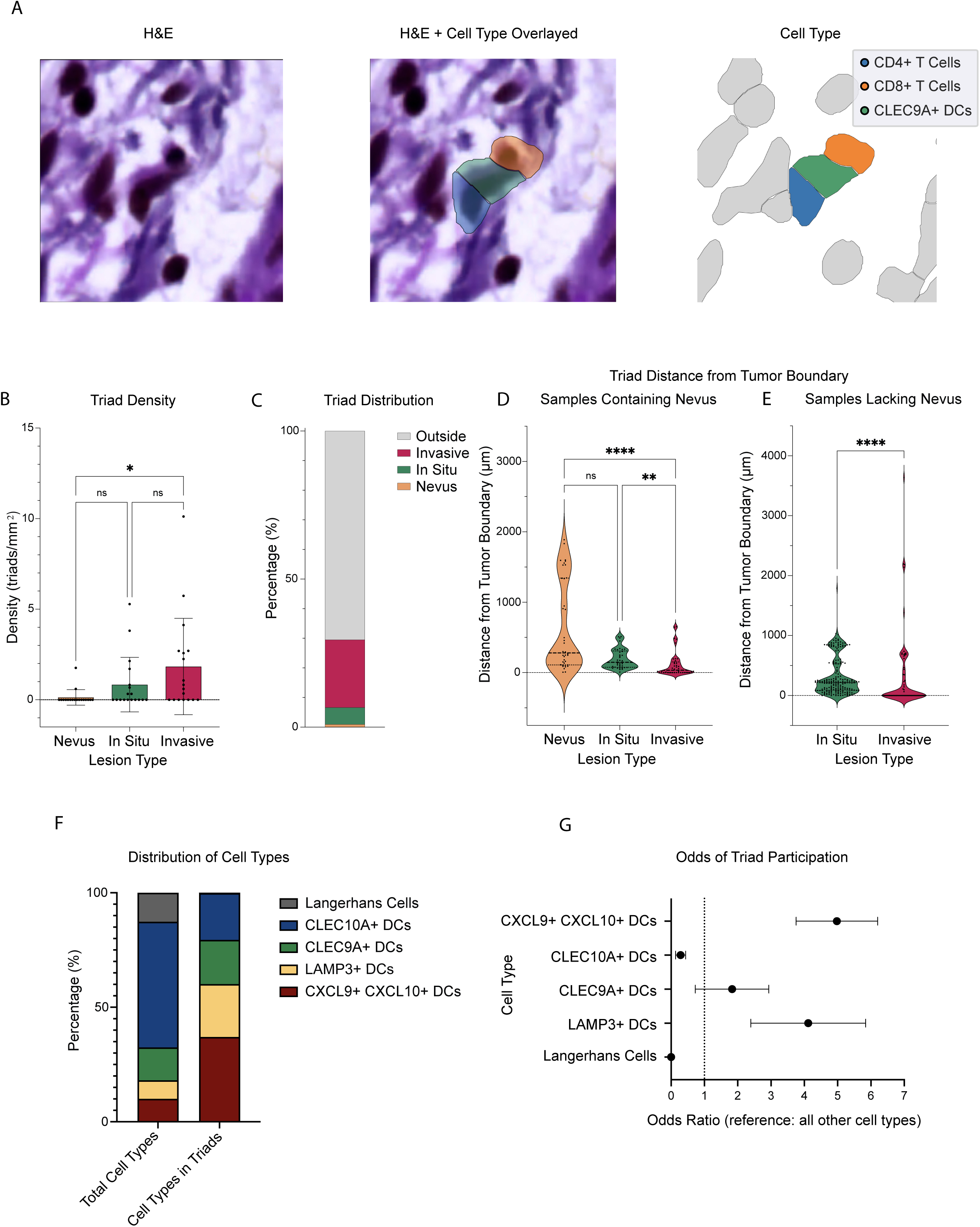
Immune triads are increased in and around regions of invasive melanoma. (A) Representative image of a CD4+ T cell, CD8+ T cell, and DC forming a triad. Panels show H&E (left), H&E overlaid with Xenium-identified CD8+ T cell, CD4+ T cell and CLEC9A+ DC cell outlines (middle), and the same CD8+ T cell, CD4+ T cell and CLEC9A+ DC showing Xenium-identified cell outlines alone (right). (B) Bar graph showing density of triads by region type. (C) Stacked bar graph showing the percentage of total triads in each region type and outside dermatopathologist-annotated regions. (D, E) Violin plot showing the distance from each triad located outside the annotated tumor regions to the nearest boundary of each region type, for samples containing nevus (D) or lacking nevus (E). (B, D-E) Statistical significance based on Kruskal-Wallis test, \*\**p*<0.01, \*\*\*\**p*<0.0001. (F) Stacked bar graph of distributions of CLEC10A+ DCs, CLEC9A+ DCs, LAMP3+ DCs, CXCL9+ CXCL10+ DCs, and Langerhans cells (LCs) considering all DCs and LCs in the samples (left) and considering only DCs and LCs participating in triads (right). (G) Odds ratio plots showing odds of DC subtypes and LCs participating in triads with participation of all other DC subtypes and LCs as a reference. An OR > 1 indicates higher odds of the indicated cell type participating in a triad. Error bars show 95 percent confidence interval.

Additionally, in line with prior reports that triads can also be found at the tumor edge or in tumor-adjacent stroma, we observed that many triads in our samples (70.5%) were found outside of annotated tumor regions (**Fig 4C**). We next measured the distance between each triad and the nearest melanoma region corresponding to each stage of disease (nevus, in situ and invasive). We found that triads external to annotated regions were located significantly closer to invasive melanoma than to nevi or to regions of in situ disease (**Fig. 4D**). This was consistent across specimens where all three disease stages were present as well as specimens which only contained in situ and invasive regions (**Fig 4E**). Additionally, while not significant, triads trended toward being closer to in situ disease than to nevi. These findings suggest that active immune cell engagement increases as melanoma progresses, both within invasive melanoma regions and in their surrounding spatial vicinity.

We found that multiple types of dendritic cells participated in triads. LAMP3+ DCs, CXCL9+ CXCL10+ DCs, CLEC9A+ DCs, and CLEC10A+ DCs were all found in triads, but we observed their composition differed from that of the total DC population (**Fig. 4F**). LAMP3+ DCs and CXCL9+ CXCL10+ DCs were enriched in triads whereas CLEC10A+ DCs were less likely to participate. Consistent with this, odd ratios analysis showed that LAMP3+ DCs and CXCL9+ CXCL10+ DCs had odds ratios greater than 1 (LAMP3+ DCs OR: 3.034; CXCL9+ CXCL10+ DCs OR: 3.574) (**Fig. 4G**), indicating greater odds of triad participation relative to all other dendritic cell types. CLEC10A+ DCs had an odds ratio less than 1 (OR: 0.173), indicating lower odds of participating in triads (**Fig. 4G**). CLEC9A+ DCs showed no significant enrichment or depletion. These findings are consistent with prior literature: cDC1s (CLEC9A+) have been reported to be major participants in triads, but when activated they transcriptionally downregulate traditional cDC1 markers in favor of an “activated” transcriptional profile (e.g., LAMP3+)(24). As such, the LAMP3+ and CXCL9+CXCL10+ populations may reflect engaged cDC1s. By contrast, our data are consistent with CLEC10A+ cDC2s being less likely to participate in T cell interactions, but we observe they do sometimes form triads.

### Langerhans cells do not participate in immune triads

Langerhans cells (LCs) are a specialized, skin-resident immune population that play an important role in immune surveillance in the skin. Although macrophage-like in origin, Langerhans cells are unique among tissue-resident macrophages in that they can uptake antigens and traffic to region lymph nodes to prime a T cell response, a function typically played by dendritic cells. Given the well-known DC-like function of Langerhans cells, we asked whether LCs also participate in immune triads with CD4+ and CD8+ T cells in or around melanoma. Strikingly, we found they did not. We observed only two instances of Langerhans-CD4-CD8 triads across our 15 specimens (**Fig. 5A**), in contrast to 239 triads formed with DCs. The fraction of LCs participating in triads was significantly lower than the fraction of DCs participating in triads (**Fig. 5B**), and this was further supported by the odds ratio analysis which showed that LCs had very low odds of participating in triads (**Fig. 4G**).

**Figure 5.**
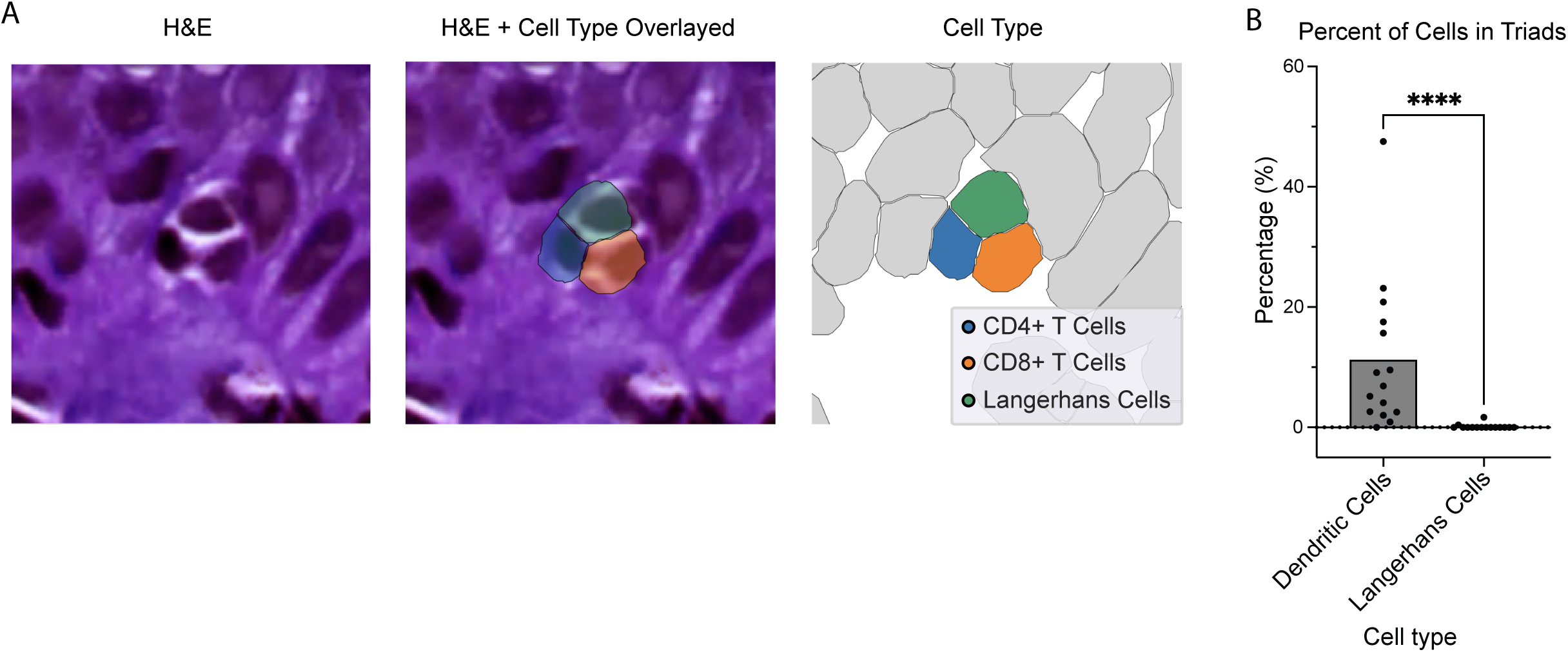
Langerhans cells rarely participate in immune triads. (A) H&E (left) and cell boundary overlay of a CD8+ T cell, CD4+ T cell and Langerhans cell (middle, right) showing a rare example of a triad including a Langerhans cell. (B) Bar graph showing percent of all DCs and of all Langerhans cells participating in a triad. Each point represents a sample. Statistical significance was evaluated by Mann-Whitney test, \*\*\*\**p*<0.0001.

### Triad abundance associates with T cell and DC density as well as T cell-DC clustering

Given the importance of triads in effective immune control of tumors, we next asked if we could identify features of regions that corresponded with the emergence of triads. Spearman correlation analysis showed that triad abundance was positively associated with the density of both CD4+ and CD8+ T cells (**Fig. 6A-B**, CD4+ T cells: ρ = 0.66, p < 0.0001; CD4+ T cells: ρ = 0.68, p < 0.0001) as well as with the density of DCs (**Fig. 6C**, ρ = 0.48, p < 0.0001) across regions of all stages of progression, suggesting perhaps not surprisingly these structures are more likely to form in regions where their constituent cells are more abundant. We next assessed spatial clustering between T cells and dendritic cells using the cross-type Ripley’s K function, which yielded a cross-type RKC score and Monte Carlo p-value. This metric quantifies whether T cells and DCs occur near each other more or less frequently than expected based on simulated null spatial distributions over the 7-35μm range. We found in both in situ and invasive regions that regions in which no significant T cell-DC clustering was observed almost universally failed to contain triads, with triads found in 0 of 5 in situ and 1 of 5 invasive regions where there was no significant T cell-DC clustering (**Fig. 6D-E**). By contrast, triads were significantly more likely to form in in situ or invasive regions where T cell-DC clustering was present (*p* = 0.02 for in situ regions and *p* = 0.04 for invasive regions, measured by chi-squared test), with triads found in 6 of 10 (60%) in situ and 16 of 23 (70%) invasive regions that met this criterion (**Fig. 6D-E**). Thus, while some degree of larger-scale spatial clustering between T cells and DCs appears to be required for the formation of triads, the presence of this clustering alone was not sufficient to definitively predict where triads would successfully form.

**Figure 6.**
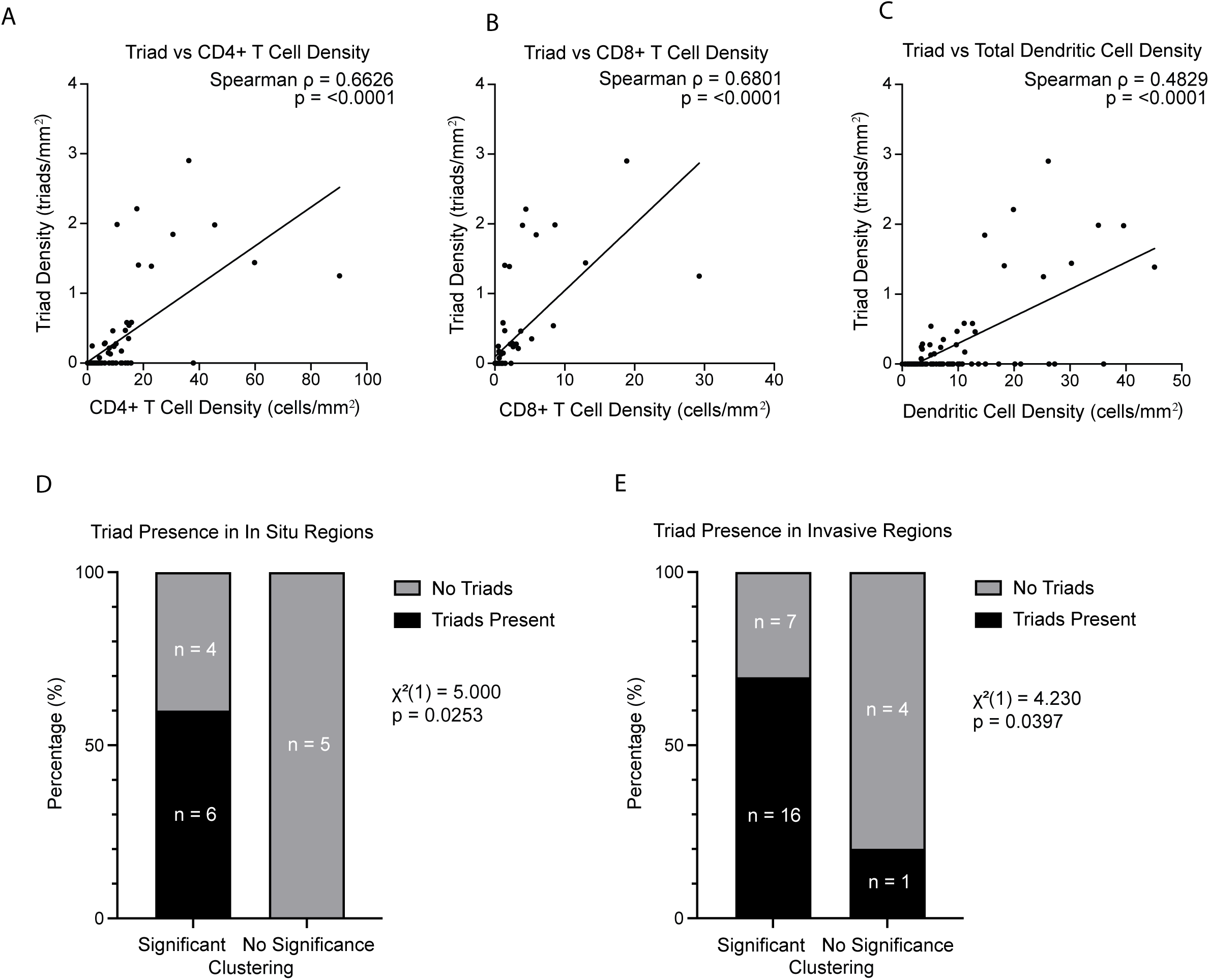
Spatial correlates of immune triad formation. (A-C) Triad density was plotted against density of CD4+ T cells, CD8+ T cells, and total dendritic cell density, and relationships were evaluated by Spearman correlation. (D, E) Stacked bar plots showing regions where T cell-DC clustering was significant (“significant”) or non-significant (“No significance”) based on Monte Carlo analysis (see Methods) and fraction of regions from each category which contained triads. Numbers within bars indicate the number of dermatopathologist-annotated regions in each category. Samples with significant T cell-DC clustering were more likely to contain triads than samples without significant clustering based on a chi-squared test.

## Discussion

We used a cohort of 15 melanoma samples containing multiple stages of progression within the same specimen to investigate the density and spatial patterns of T cell and DC infiltration across melanoma disease stages. Across melanoma progression, T cell and DC density were both significantly higher in invasive melanoma than in in situ disease and nevi and remained so when nevi and invasive lesions were matched for depth. In contrast, neither T cell clustering nor DC clustering increased with progression, as both cell types nearly always exhibited non-random clustering when invading into melanoma lesions. We additionally examined the presence of interacting multi-cell type “triads” comprising a DC, CD4+ T cell and CD8+ T cell, and found these structures were located closer to or inside regions of invasive disease relative to in situ disease or nevi. As triads are critical functional units that enhance CD8+ T cell cytotoxicity (21), this finding suggests that, beyond the mere increased numbers of immune cells present, the immune system is also more actively engaged both in and near invasive disease.

The depth-matched comparison addresses an important ambiguity in interpreting T cell infiltration and TIL infiltration more generally. Because invasive disease is deeper than Stage 0 in situ disease, increased infiltration could in principle reflect the anatomical accessibility of the dermis rather than malignancy. By using benign nevi that occupy the dermis at depths comparable to invasive lesions, we show that depth-matched benign lesions do not recruit T cells or DCs to the same extent, nor do DC-T cell triads form near them with the same prevalence, so anatomical location is insufficient to explain the enrichment. This points instead toward a driver intrinsic to the malignant cells, specifically Stage I invasive melanoma cells, that leads to enhanced immune recognition and activity – a hypothesis these data generate but do not directly test, and one that motivates future mechanistic work.

Given the importance of T cell-DC triads in immunotherapy response (21–23), we also asked what features of a lesion might predict the formation of these triads. We found that triad density was significantly correlated with the density of each constituent cell type, namely CD8+ T cells, CD4+ T cells, and DCs. We also found that multiple subsets of DCs could participate in triads, with CLEC9A+ cDC1s, CLEC10A+ cDC2s, activated LAMP3+ DCs and CXCL9+CXCL10+ DCs all found engaged in these structures. However, LAMP3+ and CXCL9+CXCL10+ DCs – both of which reflect an activated transcriptional profile – were statistically enriched in triads, while CLEC10A+ DCs were found to be less likely to participate. The engagement of DCs expressing very high levels of T cell-attracting chemokines *CXCL9* and *CXCL10* is consistent with prior reports of the importance of these signals in the formation of multicellular T cell structures (19,30). By contrast, Langerhans cells, despite being known to have DC-like functions, were rarely found to participate in triads. This aligns with a prior report suggesting Langerhans cells fail to detect early melanomas (31), and emphasizes the specific importance of conventional DC populations in melanoma immunity.

Several limitations temper these conclusions. Sample size within each progression stage was modest, and pathologist-annotated nevi regions were fewer than those for in situ or invasive disease. The study is also cross-sectional rather than longitudinal. Xenium captures immune cell spatial organization only at the time of collection, so we cannot establish the temporal order in which spatial relationships arise. Additionally, we do not have follow-up data on these patients enabling us to correlate the patterns we see with outcomes. For early-stage melanomas, such as these, long-term follow-up would be needed to accurately capture patient progression, which can occur decades after the initial melanoma is removed. Future work could carry out such correlative work with progression and survival outcomes or could apply these methods to treatment-response cohorts to relate immune cell spatial organization to therapy outcomes. Additionally, integrating Xenium data with single-cell RNAseq data may allow for finer cell-state resolution as well as comparisons of the transcriptional profiles of T cells or DCs which are engaged in triads or larger scale clustering to determine drivers of these organizational patterns.

Our findings provide strong motivation for future work examining the melanoma cell-intrinsic changes that occur between Stage 0 and Stage I disease to identify the mechanisms by which immune cells gain a dramatically increased ability to detect and engage with melanoma cells upon progression.

## Methods

### Sample selection, preparation and Xenium workflow

Formalin-fixed paraffin-embedded (FFPE) primary melanoma specimens with or without nevus precursors were identified from the University of Utah Dermatopathology archives and sectioned per 10x Genomics manufacturer recommendations. Use of human specimens for this work was approved under IRB #00172619. Xenium 5K spatial transcriptomics and subsequent Hematoxylin & Eosin (H&E) staining was performed at the South Australian Genomics Centre. Post-transcriptomic H&E images were reviewed by a board-certified dermatopathologist to assign regions of nevus, melanoma in situ, and invasive melanoma components. Coordinates for these regions, as well as the granular zone within the epidermis, were drawn in Xenium Explorer v3.0 and exported for subsequent spatial analyses.

### Xenium data processing and cell type identification

Xenium data were analyzed in python using scanpy (v1.11.0)(32), squidpy (v1.6.4)(33), and spatialdata (v0.3.0)(34) tools. Individual cells were filtered to remove cells with <100 transcripts. Leiden clustering was performed using leidenalg (v0.9.1)(35) on all cells merged across all samples to identify major cell types using canonical cell type markers. Immune cells identified in this analysis were subsequently re-clustered, and immune cell subsets identified and labeled using well-established cell type-specific markers.

### Region Import from Xenium Explorer

To convert the coordinate regions drawn in Xenium Explorer v.3.0 into Python compatible spatial objects the Xenium selection output was processed using the function xenium_explorer_selection from the spatialdata_io python package (v0.2.0)(34). Values were multiplied by 0.2125 to convert to original units (um), per spatialdata_io documentation. The resulting coordinates were transformed into valid polygon geometries using the shapely python package (v 2.1.2)(36) and exported as a GeoJSON file. The polygon files were read using GeoPandas (v1.1.2)(37) and combined into a single GeoDataFrame. To ensure consistency, all geometries were standardized to a common coordinate reference system (CRS; EPSG:3857). A spatial join was performed using GeoPandas (v 1.1.2) with geometric operations handled by Shapely (v2.1.2) to merge polygon geometries with immune cell locations. Immune cell locations were represented as a point located at the x,y cell centroid position, extracted from the xenium dataset using the get centroid function from the spatialdata package (v0.3.0).

### Density calculations

The immune cell densities were calculated within the defined polygon regions. To determine the number of immune cells in each region, Polygon area was calculated from the polygon geometry using the shapely package (v2.1.2), and cell density was calculated as

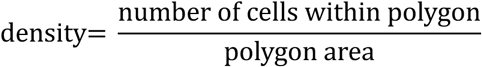

Density values were calculated separately for each immune cell type and each polygon region. These spatial analyses and density calculations were done in Python (v3.11.9) using the GeoPandas (v1.1.2), Pandas (v2.3.3) (38), and NumPy (v2.4.2) (39) libraries.

### Ripley’s K function

The Ripley’s K function (40) was used to determine spatial clustering of immune cells within melanoma tumor regions. For a point pattern containing *n* points within an observation window of area *A*, the K function is defined as

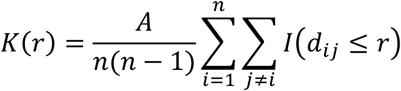

where *d_i_*_j_ is the Euclidean distance from points *i* and *j*. The Ripley’s K function is the expected number of additional points within distance r of a typical point normalized by the global point density. Under complete spatial randomness in two dimensions the expected value is *K*(*r*) = *πr*^2^ (41).This statistic was transformed to its variant the L function defined as

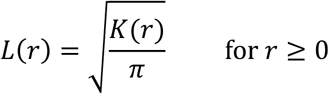

expressed as *L*(*r*) − *r* where positive values indicate clustering and negative values indicate dispersion relative to complete spatial randomness.

The Ripley’s K functions were computed by using the R spatstat package (v3.5.2)(42), via rpy2 (v3.6.4)(43). with Ripley’s isotropic edge correction to account for boundaries of the observation window. The Ripley’s K function with Ripley isotropic edge correction is defined as

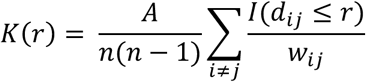

where *w_i_*_j_ is the edge correction weight, accounting for missing neighboring cells near boundaries. For intra-type analyses the K function, Kest, was computed for each cell type separately using the analysis window from the polygon. For cross-type analysis the cross-K function, Kcross, was computed for ordered pairs using a marked planar point pattern. The Ripley’s K cross function is defined as

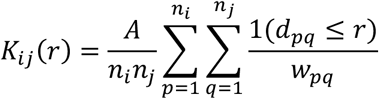

where *d_pq_* is the distance between two cells of different types. Ripley’s K analysis was only performed if 5 or more cells per cell type were present; in some cases, more stringent cut-offs were used (>30 cells for analyses in Supp. Fig. 2). Monte Carlo simulations (*n_sim_* = 1,000) were used to generate pointwise Ripley’s K simulation envelopes under the null model. Ripley’s K analyses were performed in R (v4.5.1)(44) using spatstat.geom (v3.6.0) and spatstat.explore (v3.5.2), accessed from Python through rpy2 (v3.6.4).

### Ripley’s K score (RKC score)

To summarize T cell-DC spatial association, the RKC score was defined as the integral of the Ripley’s K intra and cross L function described above:

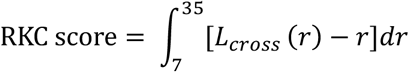

The integral was estimated numerically using the trapezoidal rule in NumPy (v 2.4.2). A RKC score greater than zero showed spatial T cell, DC, and T cell-DC clustering, where a RKC score below zero indicated spatial dispersion. The RKC score was calculated separately for each tissue polygon and ordered cell-type pair.

### Monte Carlo Simulations and p-values

Statistical significance was evaluated using 1,000 Monte Carlo simulations under complete spatial randomness. Each simulated pattern was generated within the same spatial window and preserved the same cell densities. For intra-type analyses, the observed number of cells belonging to the cell type being evaluated is preserved. The univariate K function was calculated for each simulated pattern. For cross-type analysis cell type labels were assigned independently according to the relative frequencies of T cells and dendritic cells. The cross-type K function was calculated for every simulated pattern. For both analyses, the resulting K function was integrated across 7-35μm to produce a simulated RKC score. The 1,000 simulated scores formed the null distribution which the observe scored was compared against. The upper-tail Monte Carlo probability was calculated as:

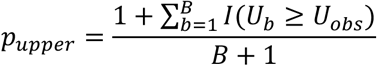

The lower tail Monte Carlo probability was calculated as:

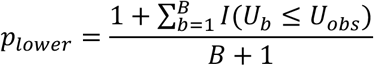

Where *B* = 1,000, *U_b_* is the integrated RKC score from simulation *b*, *U_obs_* is the observed RKC score, *I*(·) is an indicator function that equals 1 when the specified condition is satisfied and 0 otherwise.

For both intra and cross-type analysis the two-sided Monte Carlo p-value was calculated by doubling the smaller of the two tail probabilities. The addition of 1 to the numerator and denominator provides the Monte Carlo plus-one correction and prevents a p-value of zero. With 1,000 simulations the smallest possible p-value was 0.002. A two-sided p-value below 0.05 was considered statistically significant. For intra-type analyses, a significant positive RKC score indicated clustering among cells of the same type, whereas a significant negative score indicated intra-type spatial dispersion. For cross-type analyses, a significant positive RKC score indicated T cell-DC clustering, whereas a significant negative score indicated spatial dispersion between the two cell types.

### Depth

Granular zone annotations, created in Xenium Explorer v3.0 as described above, were imported using spatialdata-io v0.2.0 and converted to a LineString, and the distance between each cell centroid and the granular zone was calculated using Shapely v2.0.7. The depth of each region was calculated as the average of the depth of all cells (immune and non-immune) in the region.

### Grid analysis

A uniform square grid (160 x 160 µm) was designed for linear regressions and overlaid across the tissue regions. Grid boundaries were defined by the minimum bounding box that encompassed the annotated regions. For each individual grid square, fractional area occupied by each lesion region (nevus, in situ, invasive) was computed by intersecting the grid square with region polygons and summing the areas per region type.

### Linear Regression and Odds Ratios

To evaluate association between lesion region and immune cell abundance linear regression analysis was performed at the grid level. For each immune cell type, the cell counts per grid were modeled as a function of the fractional area of each lesion type (nevus, in situ, invasive). Linear regression models were fit to estimate the relationship between region fraction and immune cell count. Odds ratios were calculated by comparing immune cell counts between lesion regions using contingency tables with invasive lesions used as the reference category. Therefore, an odds ratio less than 1 indicates lower odds in the comparative lesion relative to invasive lesions, while an odds ratio greater than 1 indicates higher odds in the comparative lesion.

### Identification of DC-T cell triads

To identify spatial immune cell triads, Xenium data was subset to include LAMP3+, CLEC9A+, CLEC10A+, and CXCL9+ CXCL10+ dendritic cells, CD4+ T cells and CD8+ T cells. The cell centroid coordinates were used to construct a spatial proximity graph in which cells separated by ≤10.625 µm (≤50 pixels) were connected by an edge. Connected components were identified using the connected_components function from the NetworkX Python package (v3.6.1)(45). Only connected components containing at least three cells were retained. A DC-T cell triad was required to contain at least one dendritic cell (LAMP3+, CLEC9A+, CLEC10A+, or CXCL9+ CXCL10+ DCs), one CD4+ T cell and one CD8+ T cell. Triads were assigned to pathologic regions (nevus, in situ, and invasive) according to the location of the cluster centroid relative to manually annotated tissue polygons. A point in polygon approach using Shapely (v2.1.2) geometry operations were used to determine region location. For each DC in an identified triad, Euclidian distances were calculated from the DC centroid to the nearest polygon representing each region type using the shortest point to polygon distance. This same method was applied for analysis of triads containing Langerhans cells.

### Statistical analysis

Statistical analysis was performed using GraphPad Prism (v11.0.0). Comparisons between two independent groups were performed using two-sided Mann-Whitney U tests. Comparisons among three independent groups were performed using Kruskal-Wallis tests, followed by Dunn’s multiple-comparisons test. Associations between continuous variable were assessed using two-sided Spearman rank correlations, with Spearman’s correlation coefficient (ρ) reported. Associations between categorial variable were evaluated using chi-squared tests. Significance of Ripley’s K-based clustering was evaluated by Monte Carlo methods as described above. P-values below 0.05 were considered statistically significant.

## Data & Code Availability

All data and code required to reproduce these analyses are available from the corresponding authors upon reasonable request.

## Acknowledgments

This work was supported by the Huntsman Cancer Institute Cancer Center Support Grant P30CA040214; NIH R37CA304560 (to M.Q.R.); NIH R01CA228196, R01AR041836 and the J.L. Mayberry endowment (to K.J.G.); the University of Utah Department of Dermatology and the Huntsman Cancer Foundation. M.S.S. was supported by a National Health and Medical Research Investigator Award (2033440).

---

**Supp. Figure 1.**
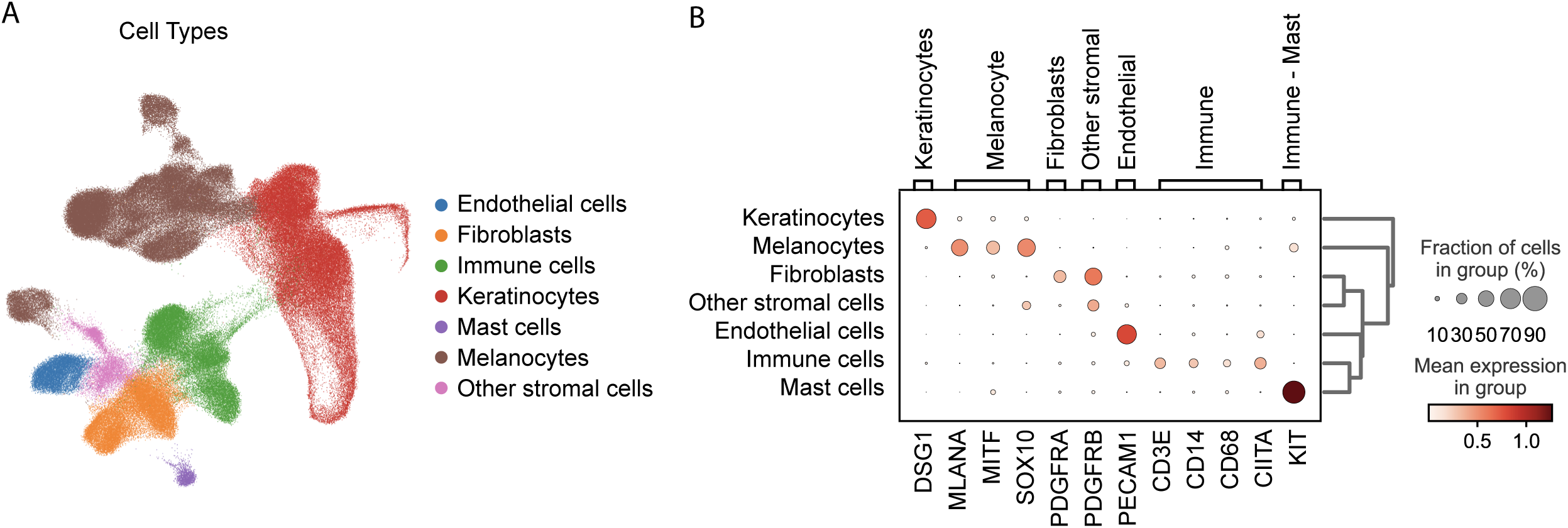
Identification of cell subsets in Xenium dataset. (A) UMAP showing all major cell types annotated in the Xenium dataset. (B) Dot plot showing expression of canonical cell type markers across annotated populations.

**Supp. Figure 2.**
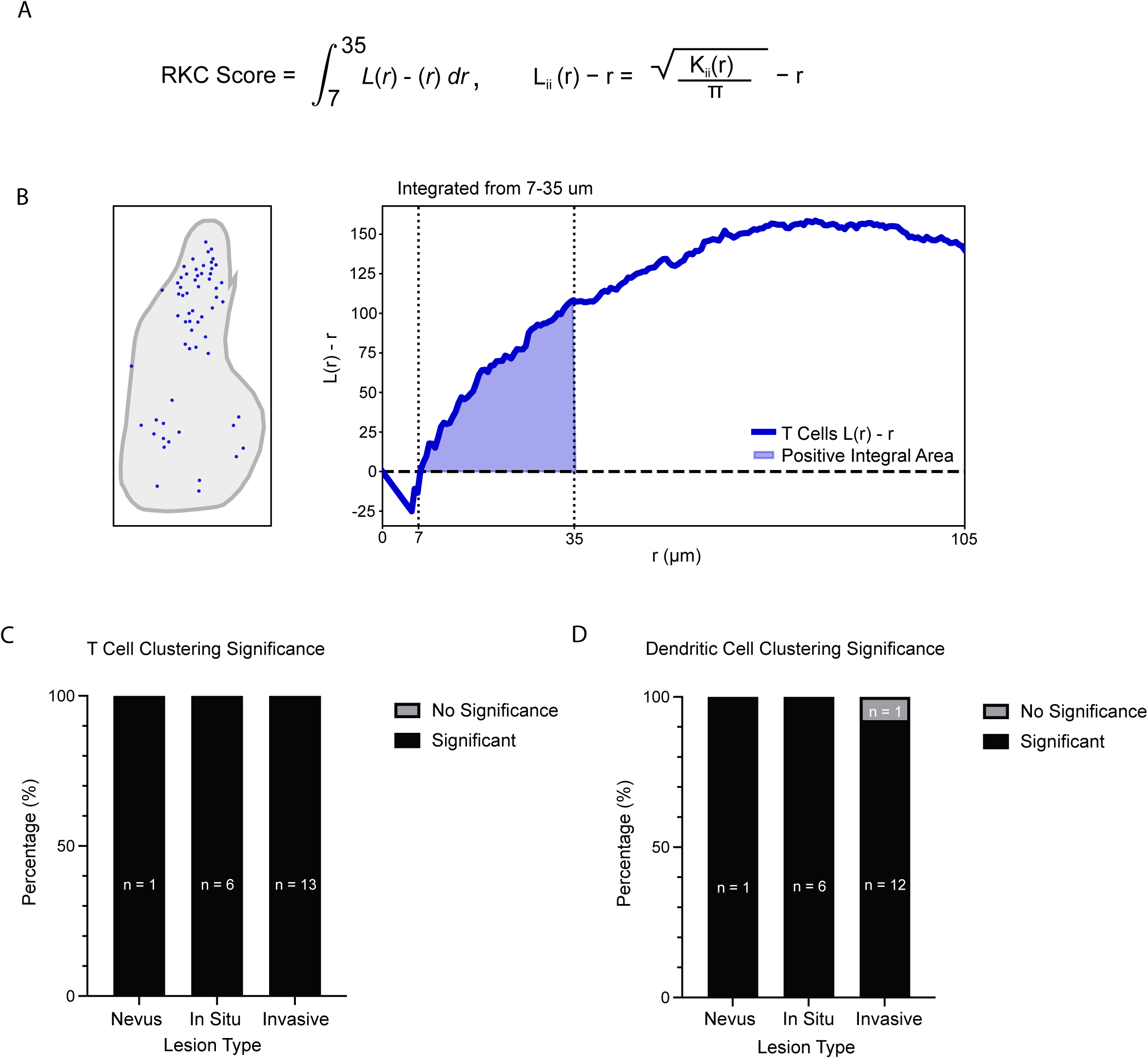
RKC score-based analysis of immune cell clustering. (A) Definition of the Ripley’s K Clustering (RKC) score. RKC score was calculated by integrating L(r)-r from 7-35μm. (B) A representative lesion region with positive RKC score (reflective of clustering) and corresponding L(r)-r curve for T cells. The RKC score is the shaded area under the curve from 7-35μm. A positive area reflects spatial clustering (shown in this example), while a negative area would reflect dispersion (not shown). (C, D) Bar graphs showing percentage of nevus, in situ, and invasive regions with significant T cell (C) or dendritic cell (D) clustering. Only regions with ≥30 cells were included for analysis. Numbers within bars indicate the number of regions. Significance was based on Monte Carlo simulations (see Methods).

